# Bowel dysmotility and enteric neuron degeneration in lysosomal storage disease mice is prevented by gene therapy

**DOI:** 10.1101/2023.05.26.542524

**Authors:** Matthew J. Jansen, Letitia L. Williams, Sophie H. Wang, Elizabeth M. Eultgen, Keigo Takahashi, Hemanth R. Nelvagal, Jaiprakash Sharma, Marco Sardiello, Brian J. DeBosch, Jessica B. Anderson, Sophie E. Sax, Christina M. Wright, Takako Makita, John R. Grider, Mark S Sands, Robert O. Heuckeroth, Jonathan D. Cooper

**Author notes:** Author for correspondence: Jonathan D. Cooper, Departments of Pediatrics, Genetics and Neurology, Washington University in St. Louis, School of Medicine, 660 S Euclid Ave, St Louis, MO 63110, USA. Authors share co-first authorship (Matthew J. Jansen, Letitia L. Williams, Sophie H. Wang). Co-senior authors (Robert O. Heuckeroth, Jonathan D. Cooper). **Authors’ contribution:** JDC, ROH, and MSS conceived and designed the study and obtained funding; MJJ, JDC, and SHW carried out the bowel whole mount pathology experiments, including immunostaining and quantitative analyses; MJJ prepared all the figures with JDC; LLW performed bowel transit studies, with initial input from KT; EME managed the mouse colonies, generated the mice for these studies and performed all genotyping; HRN and KT performed brain histology, staining and analyses with MJJ; JS and MS constructed the AAV9- CLN2 vector used in these studies; BJD assisted in interpreting data; JBA, SS and CMW performed the organ bath studies of bowel contractility under the supervision of ROH, with additional data analysis by JRG; TM performed the initial pilot studies with JDC and MSS; MSS performed the the neonatal vector administration; JDC, MSS and ROH supervised all studies and interpreted data; JDC and ROH wrote the manuscript with input from all the authors. All authors read and approved the final manuscript. **Data Transparency Statement**: All the data from this study will be made available to other researchers.

## Abstract

**Background and aims:** Children with neurodegenerative disease often have debilitating gastrointestinal (GI) symptoms that may be due at least in part to underappreciated involvement of neurons in the enteric nervous system (ENS), the master regulator of bowel function.

**Methods:** We investigated bowel motility in mouse models of CLN1 and CLN2 disease, neurodegenerative lysosomal storage disorders caused by deficiencies in palmitoyl protein thioesterase-1 (PPT1) and tripeptidyl peptidase-1 (TPP1), respectively. We then explored the integrity of ENS anatomy in immunostained bowel wholemount preparations from these mice. Lastly, we administered adeno-associated viral gene therapy to neonatal mice and determined if this would prevent these newly identified bowel phenotypes.

**Results:** Mouse models of CLN1 and CLN2 disease both displayed slow bowel transit *in vivo* that worsened with age. Although the ENS appeared to develop normally, there was a progressive and profound loss of myenteric plexus neurons accompanied by changes in enteric glia in adult mice. Neonatal administration of adeno-associated virus-mediated gene therapy prevented bowel transit defects and the loss of many ENS neurons.

**Conclusions:** We show that two neurodegenerative lysosomal storage diseases cause profound and progressive damage to the mouse enteric nervous system and impair bowel motility. We also provide proof-of-principle evidence that gene therapy can prevent enteric nervous system disease. This study may have general therapeutic implications for many inherited neurodegenerative disorders.

**What you need to know:** *Background and Context:* Many pediatric central nervous system disorders also have debilitating gastrointestinal symptoms. For most of these diseases, it is not known if the enteric nervous system (ENS) is also affected and to what degree ENS defects contribute to GI symptoms. To date, no attempts have been made to directly treat or prevent enteric nervous system disease via gene therapy.

*New Findings:* The enteric nervous system is severely affected in mouse models of CLN1 and CLN2 disease, profoundly neurodegenerative lysosomal storage disorders. Bowel transit defects and most of the enteric nervous system pathology can be prevented by neonatal administration of gene therapy.

*Limitations:* Information about enteric nervous system disease in human children is still lacking, and methods will need to be developed to treat the human bowel.

*Impact:* These findings identify an underappreciated effect of neurodegenerative disease upon the bowel and demonstrate that enteric nervous system degeneration can be prevented in mice. This provides a new perspective on these childhood disorders that may be applicable to many other conditions that affect the bowel.

*Lay Summary:* In children’s diseases where the brain degenerates, nerve cells in the bowel also die causing gastrointestinal problems, but this can be prevented by gene therapy.

## Introduction

Severe gastrointestinal (GI) symptoms are common in children with neurodegenerative disorders of the central nervous system (CNS)^1–6^. GI dysfunction causes malnutrition, feeding intolerance, vomiting, constipation, bowel distension, and abdominal pain that markedly impairs quality of life^7, 8^. GI symptoms are often attributed to CNS disease or medication side effects. We hypothesized that GI symptoms may be due, at least in part, to an under-recognized loss of neurons *outside* the CNS, specifically within the enteric nervous system (ENS).

The ENS is the intrinsic nervous system of the bowel that regulates most aspects of bowel function including motility, epithelial barrier activities, immune cell function, and blood flow^9–13^. To perform these roles, the ENS has about as many neurons as the spinal cord, ∼20 neuron subtypes, and several types of glia^9–13^. The ENS also interacts with extrinsic bowel innervatation^14^. ENS defects cause life-threatening human bowel motility disorders^6, 15^, and bowel-brain connections in some adult neurodegenerative diseases are well known^16^. Whether ENS neurodegeneration occurs in childhood CNS neurodegenerative disease is not yet determined.

To explore this issue we studied mouse models of profoundly neurodegenerative lysosomal storage disorders (LSDs) called neuronal ceroid lipofuscinoses (NCL or Batten disease)^17, 18^. There are >60 LSDs caused by mutations in genes critical for lysosomal function^19, 20^. Most LSDs damage the CNS and cause GI symptoms. Two of the earliest onset, most profoundly neurodegenerative NCLs, are CLN1 and CLN2 diseases^17, 18, 21–24^, caused by palmitoyl protein thioesterase 1 (PPT1)^25^ and tripeptidyl peptidase 1 (TPP1)^24^ mutations, respectively. Symptoms appear in the first (CLN1 disease) or second year of life (CLN2 disease), progress rapidly, and cause premature death. Mice modeling CLN1 (*Ppt1^-/-^*) or CLN2 disease (*Tpp1^R207X/R207X^*) display behavioral, pathological, and molecular phenotypes that recapitulate human disease manifestations^26–30^. Because CNS neurodegeneration is profound, prior treatment attempts mainly targeted the CNS disease manifestations via intracerebroventricular enzyme replacement^31^ or CNS-directed gene therapy^27, 30^. Because treated mice still die prematurely, we hypothesized that extra-CNS disease manifestations may also need treatment. We discovered that CLN1 and CLN2 mouse models have bowel dysmotility and severe progressive ENS neurodegenerative disease that can be treated via gene therapy.

## Materials and Methods

### Mice

*Ppt1^-/-^* mice^26^, *Tpp1^R207X/R207X^* mice^29^, and wild type (WT) mice were maintained separately on a congenic C57Bl/6J background at Washington University School of Medicine or Children’s Hospital of Philadelphia. Mice were provided food and water *ad libitum* under a 12hr light/dark cycle. Numbers analyzed are detailed in Figures, but all studies used balanced numbers of males and females. These studies follow ARRIVE guidelines and were performed under protocols 2018-0215 and 21-0292 approved by the Institutional Animal Care and Use Committee (IACUC) at Washington University School of Medicine in St. Louis, MO and The Children’s Hospital of Philadelphia IACUC protocol IAC 22-001041.

### Bowel transit studies

*Whole bowel transit*: Mice fasted overnight were gavage fed 6% carmine red dye (300 µl) Sigma-Aldrich) in distilled water containing 0.5% methylcellulose (Sigma-Aldrich) and placed in individual cages with white paper covering cage bottoms. Passage of red stool was evaluated at 10 min intervals. Each mouse was tested three times with ∼3 days between tests^32^.

*FITC-dextran transit*: Mice fasted overnight were gavage-fed 100 μL FITC-Dextran (10 mg/mL, 70,000 MW; MilliporeSigma) in distilled water containing 2% methylcellulose (MilliporeSigma).

After 2 hours, bowel from isoflurane-euthanized mice was divided into 15 segments (small intestine 1–10, proximal and distal cecum, colon 1–3). Segment contents were suspended in 100 µL 1X phosphate buffered saline, vortexed 15 s, and centrifuged (4000 rpm, 10 min). Supernatent fluorescence was measured (excitation 485 nm, emission at 525 nm). Weighted geometric mean (sum of segment number × (FITC fluorescence in that segment)/total FITC fluorescence)) was calculated as described^33, 34^.

*Colonic bead expulsion*: Mice were anesthetized with isoflurane (1.5 min). A glass bead (3 mm, MilliporeSigma) lubricated with sunflower seed oil (MilliporeSigma) was inserted 2 cm into colon using a custom-made 3 mm diameter rounded glass rod. Anesthesia was stopped and bead expulsion time was recorded^32^. Assay was repeated 3 times per mouse with >48 hours between procedures.

### Bowel contractility studies

Mice were euthanized via cervical dislocation. The bowel was removed and placed in warmed (37°C), oxygenated (95% O2, 5% CO2) Krebs-Ringers solution in a horizontal organ bath (Hugo Sachs Elektronik, Harvard Apparatus)^35^. Colon and small intestine motility wereSupplement evaluated as previously^35^, filming contractions over 20 minutes. For colon, 1 cm was trimmed off each end and mid-colon was evaluated. For small bowel, the most distal 2-3 cm of ileum was evaluated. Colon was tested first. Small bowel was stored in UW Belzer transplant solution at 4C for 40 minutes during colon imaging. For some experiments, after an initial 20 minute video, tetrodotoxin (TTX) at a concentration of 1 μM was pumped into the organ bath, and another 20 minute video recorded. Videos were converted to .wmv format using Movie Maker and saved at 1× and 16× speeds. In-house MATLAB (MathWorks) scripts (http://github.com/christinawright100/BowelSegmentation) were used to threshold movies, and generate kymographs. Low frequency contractions (LFCs) that propagated at least 1 cm from proximal to distal bowel were counted after initial studies confirmed these LFCs were eliminated by TTX (i.e., neurally mediated). All analyses were made blind to genotype or treatment.

### Wholemount bowel histology

Mice anesthetized (2% isoflurane) were transcardially perfused with PBS and decapitated. The first and last 8 cm of small intestine and entire colon were cut into 2 cm lengths in cold 50 mM Tris buffered saline (TBS, pH=7.6), opened along mesenteric border, pinned to Sylgard^TM^ 184 Silicone Elastomer (Dow Corning) using insect pins^32, 35, 36^, and fixed in fresh 4% paraformaldehyde (35 minutes colon, 45 minutes ileum, 1 hour duodenum). Fixed tissue was stored in TBS/0.1% sodium azide (4°C) before staining for neuronal or glial markers^32, 35, 36^.

Muscle layers (1 cm lengths) separated from mucosa and submucosa were blocked (1 hours) in TBS containing 4% Triton X-100 (TBST), 15% normal goat serum (NGS) (Jackson Immuno- Research Laboratories), then incubated in primary antibody (Supplemental Table 1) at 4°C overnight, washed (3x, TBS), and incubated in secondary antibodies (Supplemental Table 1) in 10% goat serum, 4% TBS-T for 2 hours. Bowel was mounted on Colorfrost plus (Thermo Fisher Scientific) slides, dried briefly before incubation in 1x solution TrueBlack lipofuscin autofluoresence quencher (Biotium) in 70% ethanol (5 min) and rinsing (1xTBS). Slides were coverslipped in Fluoromount-G mounting medium containing DAPI (Southern Biotech).

### Quantitative analysis of immunostained bowel

Unless otherwise specified, 10 systematically sampled regularly spaced 20× fields (each 0.42250 mm^2^) per sample were analyzed. Multi-channel images were collected on a Zeiss AxioImager Z1 microscope using StereoInvestigator (MBF Bioscience) software and exported to ImageJ (NIH). Myenteric neurons (HuC/D+ cells) and enteric glia (S100B+ cells) were manually counted in each field and averaged to determine cells/mm^2^. Areas of Iba1+ macrophages were measured for 100 cells per animal using StereoInvestigator software. All analyses were made blind to genotype or treatment.

### Neonatal intravenous gene therapy

Injections employed the same AAV9 virus expressing human PPT1 used in our CNS studies, described previously in detail^27, 38^ (*Ppt1^-/-^* mice), an equivalent virus expressing human TPP1 instead (*Cln2^R207X^* mice), or an equivalent virus expressing GFP. Viruses were packaged at University of North Carolina Vector Core. Virus injections were made into the superficial temporal vein of hypothermia-anesthetized neonatal (P1) mice, as described^39, 40^, to deliver 1.5 × 10^11^ vg/mouse. Injected mice were returned to their mothers, raised until weaning, and aged until predicted disease endstage (*Ppt1^-/-^* mice, 7 months; *Tpp1^R207X/R207X^* mice, 3.5 months) before most analyses.

### Brain processing and immunostaining

Brains removed from the same mice euthanized for bowel preparations were bisected. One hemisphere was dropfixed 48 hours in fresh 4% paraformaldehyde, then cryoprotected (30% sucrose in 50 mM TBS, pH=7.6). Forty µm coronal forebrain sections were cut using a Microm HM430 freezing microtome (Microm International) equipped with a Physitemp BFS-40MPA freezing stage (Physitemp, Clifton, NJ). Sections were collected into 96 well plates containing cryoprotectant solution, as described^27, 30, 31^.

A one-in-six series of coronal forebrain sections from each mouse were stained on slides using a modified immunofluorescence protocol^30, 31^ employing TrueBlack and stained with GFAP, CD68 and SCMAS antibodies (Table 1). To quantify AFSM accumulation and glial activation (GFAP+ astrocytes, CD68+ microglia), thresholding image analysis was performed as described^30, 31^ using slide-scanned images at 10x magnification (Zeiss Axio Scan Z1). Contours of appropriate anatomical regions were drawn and images analyzed using *Image-Pro Premier* (Media Cybernetics) and thresholds selected for foreground immunoreactivity above background.

**Table 1.**
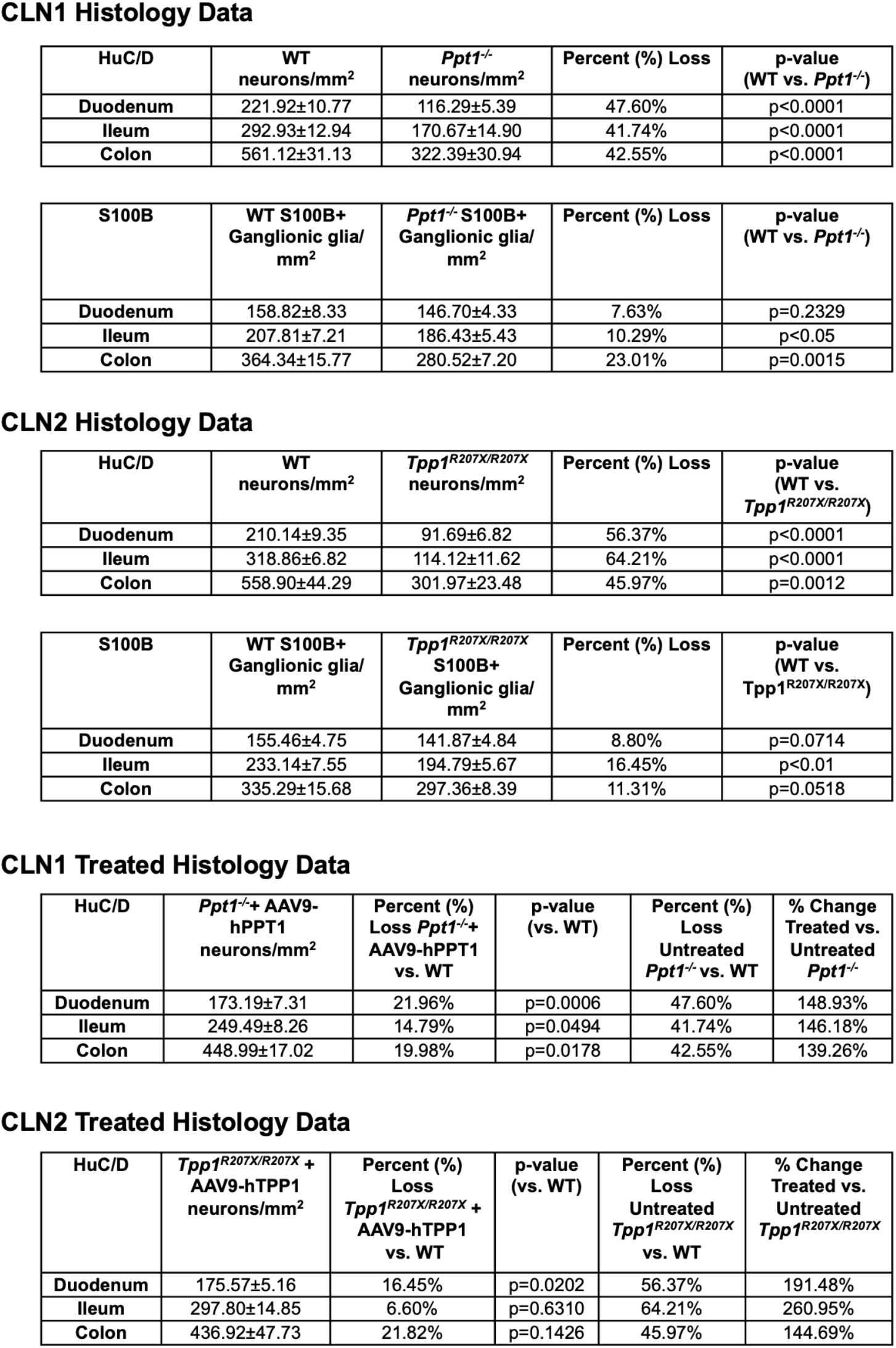
Counts of the density of HuC/D+ neurons, S100B+ enteric glia in untreated *Ppt1^-/-^* and *Tpp1^R207X/R207X^* mice, percent loss and p values. The treatment effects of AAV9-hPPT1 and AAV9-hTPP1 upon the density of HuC/D+ neurons is also listed comparing the percentage loss to both wild type (WT) and untreated mutant mice.

CLN1 Histology Data
| HuC/D | WT<br>neurons/mm<sup>2</sup> | *Ppt1*<sup>-/-</sup><br>neurons/mm<sup>2</sup> | Percent (%) Loss | p-value<br>(WT vs. *Ppt1*<sup>-/-</sup>) |
| --- | --- | --- | --- | --- |
| Duodenum | 221.92±10.77 | 116.29±5.39 | 47.60% | p<0.0001 |
| Ileum | 292.93±12.94 | 170.67±14.90 | 41.74% | p<0.0001 |
| Colon | 561.12±31.13 | 322.39±30.94 | 42.55% | p<0.0001 |

| S100B | WT S100B+<br>Ganglionic glia/<br>mm<sup>2</sup> | *Ppt1*<sup>-/-</sup> S100B+<br>Ganglionic glia/<br>mm<sup>2</sup> | Percent (%) Loss | p-value<br>(WT vs. *Ppt1*<sup>-/-</sup>) |
| --- | --- | --- | --- | --- |
| Duodenum | 158.82±8.33 | 146.70±4.33 | 7.63% | p=0.2329 |
| Ileum | 207.81±7.21 | 186.43±5.43 | 10.29% | p<0.05 |
| Colon | 364.34±15.77 | 280.52±7.20 | 23.01% | p=0.0015 |

CLN2 Histology Data
| HuC/D | WT<br>neurons/mm<sup>2</sup> | *Tpp1*<sup>R207X/R207X</sup><br>neurons/mm<sup>2</sup> | Percent (%) Loss | p-value<br>(WT vs.<br>*Tpp1*<sup>R207X/R207X</sup>) |
| --- | --- | --- | --- | --- |
| Duodenum | 210.14±9.35 | 91.69±6.82 | 56.37% | p<0.0001 |
| Ileum | 318.86±6.82 | 114.12±11.62 | 64.21% | p<0.0001 |
| Colon | 558.90±44.29 | 301.97±23.48 | 45.97% | p=0.0012 |

| S100B | WT S100B+<br>Ganglionic glia/<br>mm<sup>2</sup> | *Tpp1*<sup>R207X/R207X</sup><br>S100B+<br>Ganglionic glia/<br>mm<sup>2</sup> | Percent (%) Loss | p-value<br>(WT vs.<br>*Tpp1*<sup>R207X/R207X</sup>) |
| --- | --- | --- | --- | --- |
| Duodenum | 155.46±4.75 | 141.87±4.84 | 8.80% | p=0.0714 |
| Ileum | 233.14±7.55 | 194.79±5.67 | 16.45% | p<0.01 |
| Colon | 335.29±15.68 | 297.36±8.39 | 11.31% | p=0.0518 |

CLN1 Treated Histology Data
| HuC/D | *Ppt1*<sup>-/-</sup> + AAV9-hPPT1<br>neurons/mm<sup>2</sup> | Percent (%)<br>Loss *Ppt1*<sup>-/-</sup> +<br>AAV9-hPPT1<br>vs. WT | p-value<br>(vs. WT) | Percent (%)<br>Loss<br>Untreated<br>*Ppt1*<sup>-/-</sup> vs. WT | % Change<br>Treated vs.<br>Untreated<br>*Ppt1*<sup>-/-</sup> |
| --- | --- | --- | --- | --- | --- |
| Duodenum | 173.19±7.31 | 21.96% | p=0.0006 | 47.60% | 148.93% |
| Ileum | 249.49±8.26 | 14.79% | p=0.0494 | 41.74% | 146.18% |
| Colon | 448.99±17.02 | 19.98% | p=0.0178 | 42.55% | 139.26% |

**Table 1** Counts of the density of HuC/D+ neurons, S100B+ enteric glia in untreated *Ppt1*<sup>-/-</sup> and *Tpp1*<sup>R207X/R207X</sup> mice, percent loss and p values.
| HuC/D | *Tpp1*<sup>R207X/R207X</sup> +<br>AAV9-hTPP1<br>neurons/mm<sup>2</sup> | Percent (%)<br>Loss<br>*Tpp1*<sup>R207X/R207X</sup> +<br>AAV9-hTPP1<br>vs. WT | p-value<br>(vs. WT) | Percent (%)<br>Loss<br>Untreated<br>*Tpp1*<sup>R207X/R207X</sup><br>vs. WT | % Change<br>Treated vs.<br>Untreated<br>*Tpp1*<sup>R207X/R207X</sup> |
| --- | --- | --- | --- | --- | --- |
| Duodenum | 175.57±5.16 | 16.45% | p=0.0202 | 56.37% | 191.48% |
| Ileum | 297.80±14.85 | 6.60% | p=0.6310 | 64.21% | 260.95% |
| Colon | 436.92±47.73 | 21.82% | p=0.1426 | 45.97% | 144.69% |

### Statistical analysis

Statistical analyses were performed using GraphPad Prism version 9.1.0 for MacOS (GraphPad Software, San Diego, CA). Unpaired t-test or Mann-Whitney test were used for comparison between two groups based on data distributions. A one-way ANOVA with a post-hoc Bonferroni correction was used for comparison between three groups or more. A p-value of ≤0.05 was considered significant.

## Results

### NCL mouse models display slow bowel transit and abnormal contractility

Well-established CLN1 disease (*Ppt1^-/-^*) or CLN2 disease (*Tpp1^R207X/R207X^*) mouse models have progressive neurological decline, CNS neurodegeneration, and pathology characteristic of human NCLs^26–30, 41^. Both models die prematurely (∼8 months for *Ppt1^-/-^*, ∼4 months for *Tpp1^R207X/R207X^*). Gastrointestinal phenotypes that might contribute to early death or morbidity have not been explored. At disease endstage (7 months for *Ppt1^-/-^*and 3.5 months for *Tpp1^R207X/R207X^*), both models have small intestine and cecum distention, with hard fecal pellets in the colon (Figure 1A). Because this suggests impaired bowel motility, we evaluated whole bowel transit by gavage-feeding carmine dye. By 7 months, *Ppt1^-/-^* have significantly delayed whole bowel transit, but carmine transit was similar to WT at earlier ages (Figure 1B).

**Figure 1.**
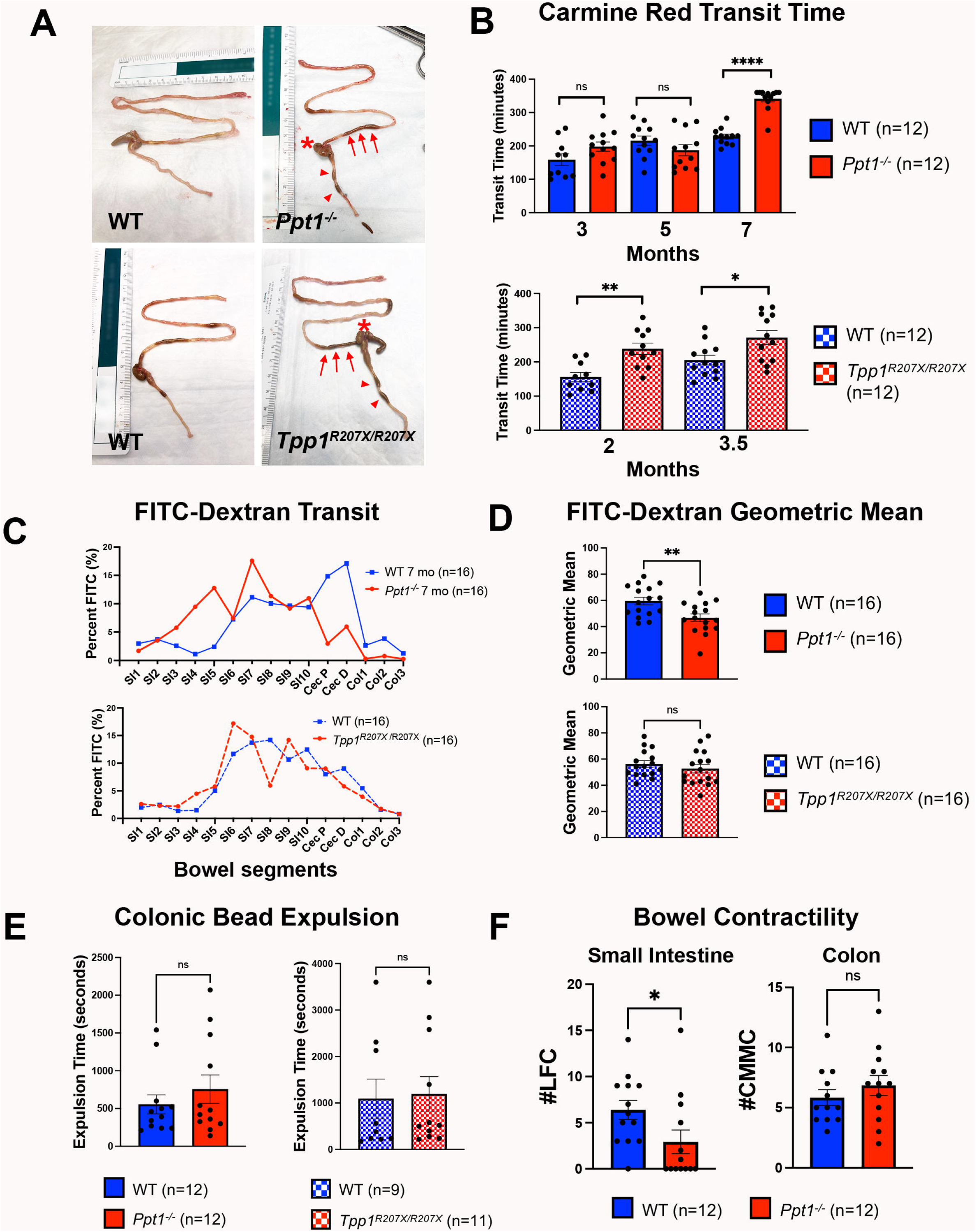
Mouse models of NCL display slow bowel transit and abnormal contractility. **(A)** CLN1 disease (*Ppt1^-/-^*) or CLN2 disease (*Tpp1^R207X/R207X^*) mice display distention of the small intestine (arrows) and cecum (*) by fecal material at disease end-stage, with hardened fecal pellets in colon (arrowheads). **(B, C, D)** Total bowel transit was assessed by measuring the time for gavaged carmine red to appear in stool (B) or determining the amount of FITC conjugated dextrans present in individual bowel segments 2 hours after gavage (C) or expressed as weighted geometic mean fluorescence (D) (SI1-SI10 = numbered segments of small intestine; Cec P=proximal cecum; Cec D=distal cecum; Col1-Col3=numbered segments of colon). **(E)** Time taken to expel a glass bead from the colon was measured at disease endstage. **(F)** Graphs of low frequency contractions per 20 minutes in small bowel or colon based on the analysis of kymographs generated from videos of bowel an oxygenated organ bath. Unpaired t- test (B, D, E, F). * p≤0.05; ** p≤0.01, **** p≤0.0001.

*Tpp1^R207X/R207X^* displayed earlier onset bowel dysmotility with delayed whole bowel carmine transit at disease midstage (2 months) and endstage (3.5 months) (Figure 1B). To evaluate transit through the stomach, small bowel, and proximal colon, we gavage-fed FITC-dextran and analyzed luminal FITC distribution 2 hours later. Seven month-old *Ppt1^-/-^* mice had much more fluorescence in proximal and mid small intestine and less in colon than age-matched WT (Figure 1C) as confirmed by weighted geometric mean (Figure 1D). Disease endstage *Tpp1^R207X/R207X^* mice also showed more FITC-dextran in small intestine and less in colon (Figure 1D), but the transit defect was less pronounced than in *Ppt1^-/-^*, and geometric means were not statistically different from controls (Figure 1C). To directly evaluate distal colon and defecatory function, we measured colonic bead expulsion times. Mean bead expulsion times were longer in *Ppt1^-/-^* than WT, but the difference was not statistically significant (Figure 1E). There was also no significant difference in colonic bead expulsion time between *Tpp1^R207X/R207X^* and WT, with high variability between animals (Figure 1E). These data show that *Ppt1^-/-^* and *Tpp1^R207X/R207X^* mice have slow transit of luminal contents *in vivo*.

To evaluate bowel motility independent of extrinsic CNS, dorsal root ganglia, sympathetic and parasympathetic innervation, we imaged *Ppt1^-/-^*bowel motility *ex vivo* in an oxygenated organ bath. Kymographs show that distal small intestine from 7-month-old *Ppt1^-/-^* mice had significantly fewer low frequency contractions (LFCs) compared to age-matched WT mice (WT 6.38±1.05 LFCs/20min; *Ppt1^-/-^* 2.92±1.28 LFCs/20min; p=0.0474, t-test), with many displaying virtually no contractions (Figure 1F, Supplemental Figure 1). These LFCs are mediated by nervous system activity since they were ablated by tetrodotoxin treatment (Supplemental Figure 1). In contrast, colon migrating motor complexes appeared similar in *Ppt1^-/-^*and WT mice (Figure 1F). Since organ bath preparations are physically isolated from extrinsic bowel innervation, these data suggest that bowel transit defects may, at least in part, be due to bowel-intrinsic defects in motility.

### Loss of myenteric plexus neurons in NCL mouse models

Because *Ppt1^-/-^* and *Tpp1^R207X/R207X^* mice have progressive degeneration of CNS neurons^27, 30, 41– 43^, we hypothesized bowel-intrinsic motility defects might result from progressive ENS damage. Alternatively, the ENS might develop abnormally. To determine if ENS appears normal before the onset of disease symptoms, we analyzed whole-mount bowel preparations from 1-month-old WT and *Ppt1^-/-^* or *Tpp1^R207X/R207X^* mice after HuC/D and S100B antibody staining but saw no overt differences between genotypes (Supplemental Figure 2A). HuC/D-positive neuron and S100B-positive glia counts revealed no differences between genotypes in any bowel region (Supplemental Figure 2B) suggesting that the ENS forms normally in *Ppt1^-/-^* and *Tpp1^R207X/R207X^* mice.

To determine if *Ppt1^-/-^* or *Tpp1^R207X/R207X^* mice undergo progressive enteric neuron damage similar to well-known CNS neurodegeneration caused by these mutations^27, 30, 41–43^, we next examined myenteric plexus at disease endstage. Immunostaining of whole-mount preparations revealed markedly fewer HuC/D-positive myenteric plexus neurons in all regions of both *Ppt1^-/-^* (Figure 2A, B) and *Tpp1^R207X/R207X^*mice (Figure 3A, B). Neurofilaments visualized with neuron-specific beta 3 tubulin antibody Tuj1 were also much less abundant in *Ppt1^-/-^* and *Tpp1^R207X/R207X^* mice at disease endstage (Figure 2D, Figure 3C). The degree of neuron loss was similar in all regions of 7 month old *Ppt1^-/-^*mice with 47.6% fewer neurons in the duodenum (p≤0.0001) and 41.74% fewer in the ileum (p≤0.0001), and 42.55% fewer neurons in the colon (p≤0.0001) of these mice (Figure 2B, Table 1). *Tpp1^R207X/R207X^* mice at disease endstage (3.5 months) had a more profound reduction in HuC/D-positive neurons in the small intestine (56.37% loss in duodenum, p≤0.0001, 64.21% in the ileum, p≤0.0001), compared to the colon (45.97% loss (p≤0.001)) (Figure 3B, Table 1). These data show that as in the CNS, profound progressive neuron loss also occurs in ENS of CLN1 disease and CLN2 disease mice.

**Figure 2.**
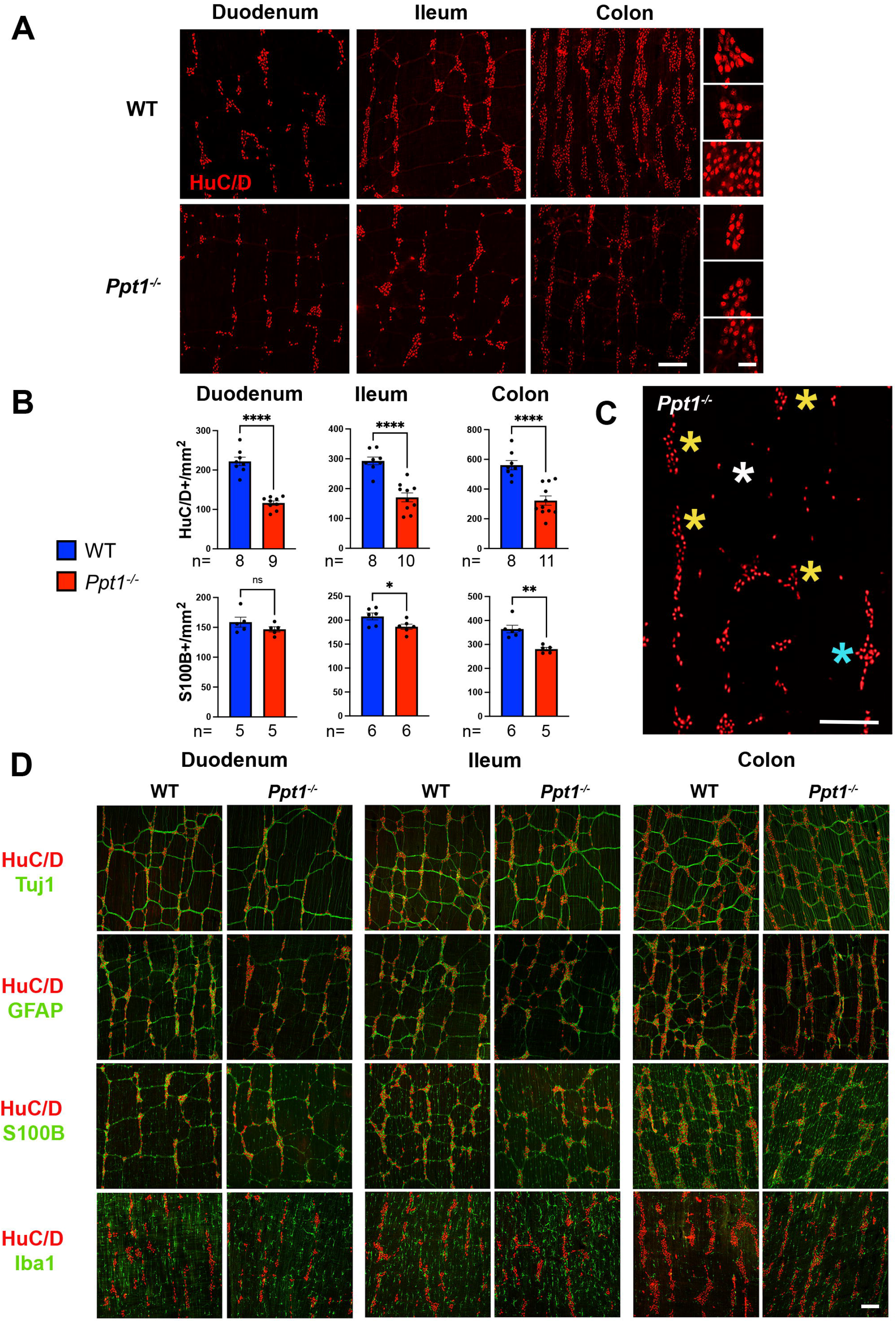
Enteric nervous system pathology in *Ppt1^-/-^* mice. **(A)** Whole-mount bowel preparations immunostained for pan-neuronal marker HuC/D (red) reveal the loss of myenteric plexus neurons and changes in their neuron morphology in the duodenum, ileum and colon of disease endstage *Ppt1^-/-^* mice vs. age matched wildtype (WT) controls at 7 months. Scale bar 200µm, 50µm inserts. **(B)** The density of HuC/D+ neurons and S100B+ enteric glia was measured in these mice at endstage. **(C)** HuC/D+ neuron loss in *Ppt1^-/-^* mice is localized, with areas of healthy appearing neurons (blue asterisk) adjacent to areas of shrunken neurons (yellow asterisk) or areas of near total neuron loss (white asterisk). Scale bar 200µm. **(D)** Merged photomicrographs showing immunostaining for HuC/D (neurons, red) and either the neuronal-specific tubulin marker Tuj1 (green), the glial marker glial fibrillary acidic protein (GFAP, green), the glial marker S100B (green), or the macrophage marker Iba1 (green) in the duodenum, ileum and colon of *Ppt1^-/-^* mice and age-matched WT controls at 7 months. (Separated channel images are included in Supplemental Figure 3). Scale bar 200µm. Unpaired t-test (B), * p≤0.05; ** p≤0.01, **** p≤0.0001.

**Figure 3.**
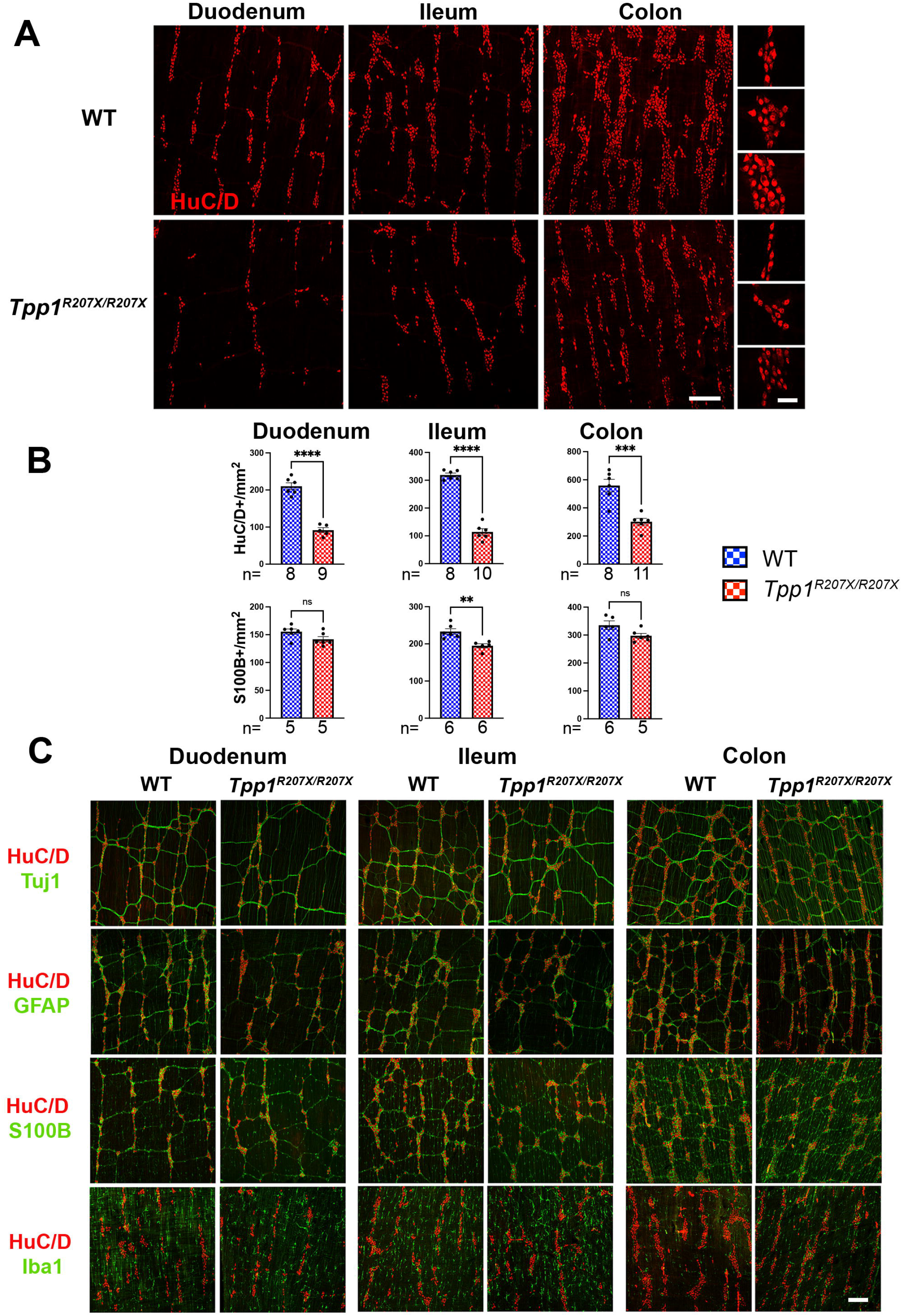
Enteric nervous system pathology in *Tpp1^R207X/R207X^* mice. **(A)** Whole-mount bowel preparations immunostained for pan-neuronal marker HuC/D (red) reveal the loss of myenteric plexus neurons and changes in their morphology in the duodenum, ileum and colon of disease endstage *Tpp1^R207X/R207X^* mice vs. age matched wildtype (WT) controls at 3.5 months. Scale bar 200 µm, 50 µm inserts. **(B)** The density of HuC/D+ neurons (B) and S100B+ enteric glia (C) was measured in the bowel of these mice at disease endstage. **(C)** Merged photomicrographs showing immunostaining for HuC/D (neurons, red) and either the neurofilament marker Tuj1 (green), glial marker glial fibrillary acidic protein (GFAP, green), the glial marker S100B (green), or the macrophage marker Iba1 (green) in the duodenum, ileum and colon of *Tpp1^R207X/R207X^* mice and age-matched WT controls at 3.5 months. (Separated channel images are included in Supplemental Figure 4). Scale bar 200µm. Unpaired t-test (B), ** p≤0.01, *** p≤0.001, **** p≤0.0001.

A striking finding was that enteric neuron loss was not evenly distributed in *Ppt1^-/-^* and *Tpp1^R207X/R207X^* mice (Figure 2A, 2C, 3A). Instead, there were patches with marked neuron loss adjacent to areas where neurons survived but had HuC/D immunoreactivity in the nucleus (Figure 2A, 2C, 3A), a pathological phenotype associated with dysfunctional enteric neurons^44^. Further away from patches of neuron loss, persisting HuC/D positive neurons appeared morphologically healthy (Figure 2C).

### PPT1 and TPP1 deficiency affects enteric glia and bowel macrophages

CNS neuron loss in CLN1 and CLN2 disease is preceded by localized astrocytosis and microglial activation most pronounced in areas where neurons are subsequently lost^30, 41–43^. To explore whether enteric neuron loss was accompanied by changes in enteric glia, *Ppt1^-/-^* and *Tpp1^R207X/R207X^* bowels were stained for glial fibrillary acidic protein (GFAP), an intermediate filament also expressed by enteric glia^37^. GFAP staining revealed areas of both increased and decreased GFAP immunoreactivity in the small intestine and colon of both models (Figure 2D, Figure 3C, Supplemental Figures 3, 4), resulting in no overall difference in GFAP immunoreactivity in any bowel region of either genotype. Nevertheless, in both *Ppt1^-/-^* and *Tpp1^R207X/R207X^* bowels areas with HuC/D-positive neuron loss consistently had less GFAP immunoreactivity (Figure 2D, Figure 3C, Suplementary Figures 3,4). Conversely, where GFAP immunoreactivity was elevated or retained, there appeared to be more HuC/D-positive neurons in both *Ppt1^-/-^* and *Tpp1^R207X/R207X^* mice (Figure 2D, Figure 3C, Suplemental Figures 3,4).

**Figure 4.**
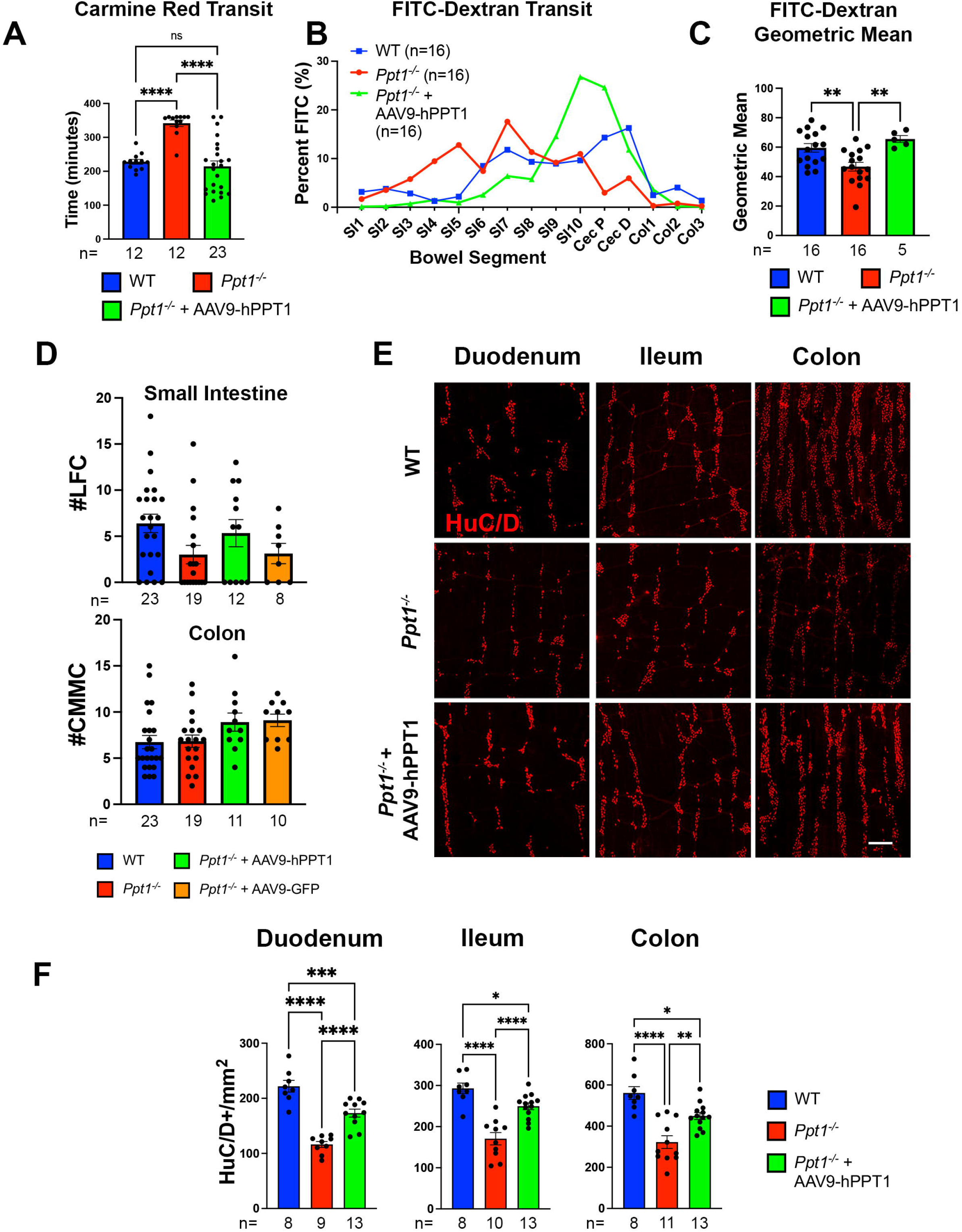
Treatment effects of gene therapy in the bowel of *Ppt1^-/-^* mice. **(A-C)** Total bowel transit was compared at 7 months in untreated *Ppt1^-/-^* mice, AAV9-hPPT1 treated *Ppt1^-/-^* mice and age-matched wildtype (WT) controls. This was done by measuring the time for gavaged carmine red to appear in stool (A) or determining the amount of FITC conjugated dextrans present in individual bowel segments 2 hours after gavage (B) or expressed as weighted geometic mean FITC fluorescence (C). (SI1-SI10 = numbered segments of small intestine; Cec P = proximal cecum; Cec D = distal cecum; Col1-Col3 = numbered segments of colon). **(D)** Bowel contractility kymographs were recorded in the small intestine and colon of mice from all treatment groups in an oxygenated organ bath and the number of low frequency contractions (LFCs) (white arrowheads on kymographs) in 20 minutes calculated. **(E)** Representative photomicrographs of whole-mount bowel preparations immunostained for the pan-neuronal marker HuC/D (red) reveal the protective effects of AAV9-hPPT1 treatment upon myenteric plexus neurons in the bowel of disease endstage AAV9-hPPT1-treated *Ppt1^-/-^* mice vs. untreated *Ppt1^-/-^* mice vs. WT controls at 7 months. Scale bar 200µm. **(F)** Counts of the density of HuC/D positive neurons reveal significantly more neurons present in all bowel regions of 7 month old AAV9-hPPT1 treated *Ppt1^-/-^* mice vs. untreated *Ppt1^-/-^* mice. One-way ANOVA with a post-hoc Bonferroni correction (A, C, F), * p≤0.05, ** p≤0.01, *** p≤0.001, **** p≤0.0001.

To determine if changes in GFAP immunoreactivity were associated with loss of enteric glia, bowels from disease end stage *Ppt1^-/-^* and *Tpp1^R207X/R207X^* mice were stained for S100B, which is co-expressed with GFAP in the soma of enteric glia, although more enteric glia express S100B (76-85%) than GFAP (35-54%)^37^. Quantitative analysis of S100B+ glial density revealed only a small reduction in ileum (10.29%, p<0.05), more loss in colon (23.01%, p<0.0015), and no significant change in duodenum (7.63% loss, p=0.2329) of *Ppt1^-/-^* mice (Figure 2B, D, Table 1). Endstage *Tpp1^R207X/R207X^* mice also had a reduction in S100B+ enteric glia density in ileum (16.45%, p<0.01), and in duodenum (8.80%, p=0.0714), and colon (11.31%, p=0.0518) that were not statistically significant (Figure 3B, Table 1).

Microglia, the only CNS myeloid cells, are involved in the pathogenesis of multiple forms of NCL^42, 43^. Intestinal macrophages are also derived from myeloid lineage and have many homeostatic roles in addition to combating pathogens^45, 46^. To determine whether myenteric neuron loss was accompanied by changes in muscularis macrophages, endstage *Ppt1^-/-^* and *Tpp1^R207X/R207X^* bowels were stained for Iba1, an established bowel macrophage marker^36^. *Ppt1^-/-^* and *Tpp1^R207X/R207X^*mice had fewer but larger Iba1-positive macrophages in all bowel regions compared to age-matched WT (Figure 2D, 3C, Supplemental Figures 3, 4). However, there was no apparent correlation between the distribution of Iba1-positive macrophages and patches where HuC/D-positive neurons were lost in either model (Figure 2D, 3C, Supplemental Figures 3, 4). Collectively, these data show changes in enteric glia and macrophages in *Ppt1^-/-^* and *Tpp1^R207X/R207X^* mice, but the relationship betwen enteric neuron loss and glial or macrophage reactivity differs from that seen in the CNS of these same models^28, 30, 41–43^.

### Adeno-associated viral gene therapy treats bowel dysfunction in NCL mice

CNS treatment strategies for CLN1 or CLN2 disease depend on overcoming PPT1- or TPP1- deficiency by supplying either recombinant enzymes^31^ or gene therapy^27–30^. To test if gene therapy could prevent newly defined functional and anatomical ENS defects in *Ppt1^-/-^* and *Tpp1^R207X/R207X^* mice, we intravenously (IV) delivered an adeno-associated viral vector (AAV9) expected to transduce enteric neurons^39, 40, 47, 48^. Because transduced neurons can secrete lysosomal enzymes that may be taken up to ‘cross correct’ neighboring cells^49^, there is no requirement to transduce all enteric neurons. Nevertheless, to confirm transduction of enteric neurons by our AAV9 vector, we first injected *Ppt1^-/-^* mice with a GFP-expressing virus (AAV9- GFP). As expected AAV9-GFP transduced a proportion of HuC/D positive enteric neurons (Supplemental Figure 6A).

To test therapeutic effects of gene therapy, neonatal *Ppt1^-/-^* and *Tpp1^R207X/R207X^* were next injected IV with previously described AAV9-vectors. Seven month-old AAV9-hPPT1-treated *Ppt1^-/-^* and 3.5-month-old AAV9-TPP1-treated *Tpp1^R207X/R207X^* had faster carmine red whole bowel transit and more FITC-dextran reaching cecum and colon than untreated controls (Figure 4A-C, 5A-C). In fact, whole bowel and FITC-dextran transit in viral vector treated mice were not significantly different from age-matched WT (Figures 4, 5). Contractility patterns of isolated small bowel and colon were also evaluated using an oxygenated organ bath. For 7-month-old *Ppt1^-/-^* mice, the mean number of propagating LFCs was reduced in untreated (3.00±1.00 LFCs/20min) and AAV9-GFP-treated mice (3.13±1.09 LFCs/20min) compared to WT (6.39±1.00 LFCs/20min) or AAV9-hPPT1-treated mice (5.33±1.47 LFCs/20min). However, there was considerable variability in LFC frequency in each group and none of these differences were statistically significant (Figure 4D).

### Adeno-associated viral gene therapy partially prevents enteric neuron loss in NCL mice

To determine if AAV9-mediated gene therapy could prevent enteric nervous system pathology, we examined bowel anatomy in AAV9-hPPT1-treated *Ppt1^-/-^* (Figure 4E) and AAV9-hPPT1- treated *Tpp1^R207X/R207X^*mice (Figure 5D) at disease end stage. In both mouse models, we saw significant treatment effects of gene therapy, at least halving the percentage of enteric neurons lost in most bowel regions (Table 1). Nevertheless, AAV9-hPPT1-treated mutant mice still had fewer neurons at disease endstage than age-matched WT (Figure 4F, 5E). However, no treatment effects were seen in AAV9-GFP treated *Ppt1^-/-^* mice (Supplemental Figure 6).

**Figure 5.**
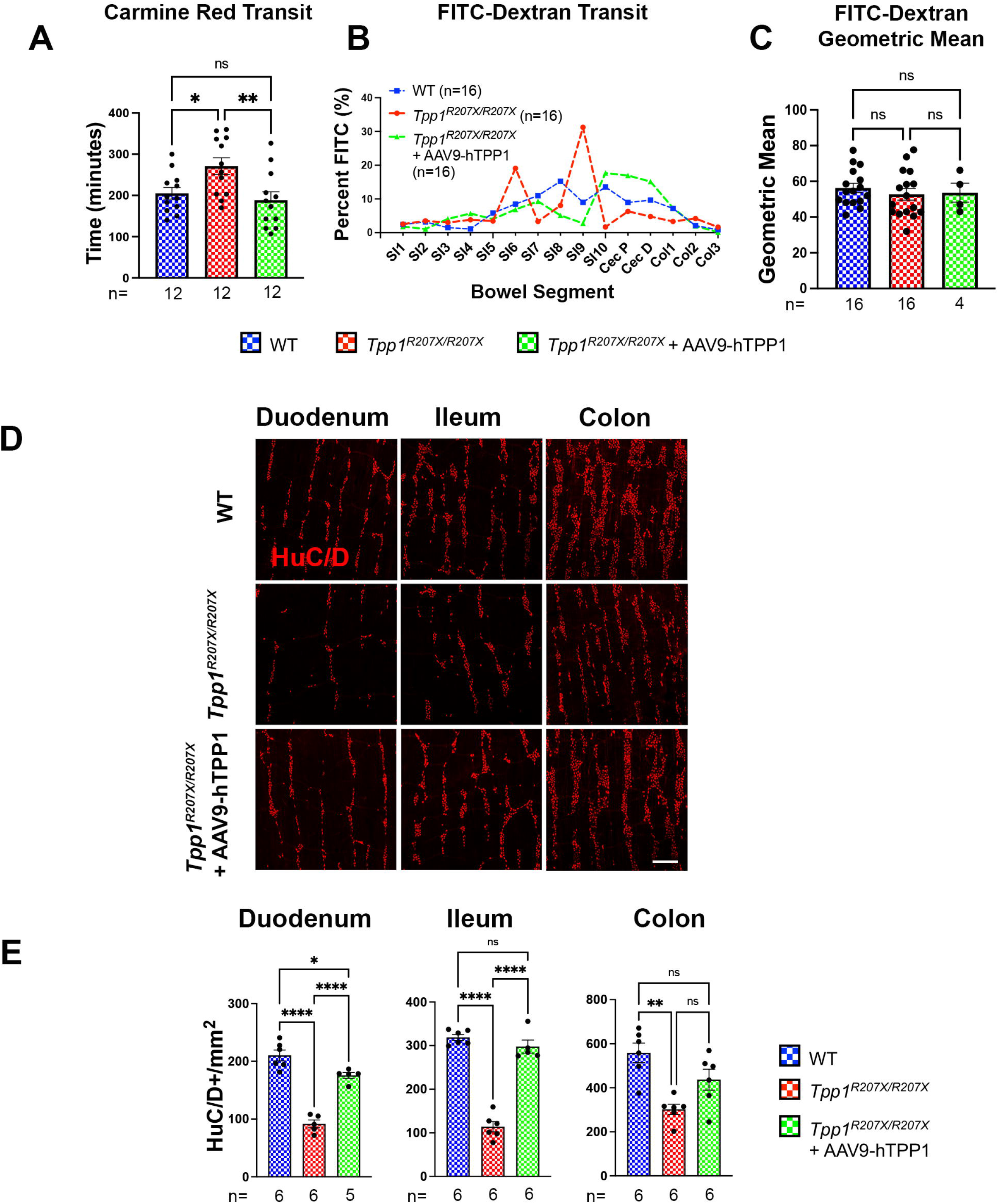
Treatment effects of gene therapy in the bowel of *Tpp1^R207X/R207X^* mice. **(A-C)** Total bowel transit was compared at 3.5 months in untreated *Tpp1^R207X/R207X^*mice, AAV9-hTPP1- treated *Tpp1^R207X/R207X^* mice, and age-matched wildtype (WT) controls. This was done by measuring the time for gavaged carmine red to appear in stool (A) or determining the amount of FITC conjugated dextrans present in individual bowel segments 2 hours after gavage (B) or expressed as weighted geometic mean FITC fluorescence (C). (SI1-SI10=numbered segments of small intestine; Cec P=proximal cecum; Cec D=distal cecum; Col1-Col3=numbered segments of colon). **(C)** Histograms displaying weighted geometic mean fluorescence revealed no difference between these values between mice of any genotype at 3.5 months. **(D)** Representative photomicrographs of whole-mount bowel preparations immunostained for the pan-neuronal marker HuC/D (red) reveal that AAV9-hTPP1 treatment prevented the loss of myenteric plexus neurons in the bowel of disease endstage AAV9-hTPP1-treated *Tpp1^R207X/R207X^* mice vs. untreated *Tpp1^R207X/R207X^* mice vs. WT at 3.5 months. Scale bar 200µm. **(E)** Counts of the density of HuC/D positive neurons reveal significantly more neurons present in the duodenum and illum, but not colon of 3.5 month old AAV9-hTPP1-treated *Tpp1^R207X/R207X^* mice compared to untreated *Tpp1^R207X/R207X^* mice. One-way ANOVA with a post-hoc Bonferroni correction (A, C, E), * p≤0.05, ** p≤0.01, **** p≤0.0001.

Collectively, these data suggest that gene therapy with enzyme-producing AAV9 vectors substantially prevents the progressive enteric neuron loss that normally occurs in CLN1 and CLN2 disease mice.

As the blood-brain barrier has not fully formed in neonatal mice^50, 51^, we also examined whether intravenous delivery of AAV9-hPPT1 or AAV9-hTPP1 treated the well-defined CNS phenotypes of *Ppt1^-/-^* and *Tpp1^R207X/R207X^*mice^27, 28, 30, 41^ (Figure 6). In both models, AAV9-mediated gene transfer reduced astrocytosis (GFAP), reduced microglial activation (CD68) and reduced storage material accumulation (SCMAS) in vulnerable regions of the thalamocortical system where neuron loss is most pronounced in untreated mice (Figure 6A-D). However, in several brain regions these parameters were not completely normalized in viral vector-treated mutant mice (Figure 6A-D). In contrast, *Ppt1^-/-^* mice treated with the control AAV9-GFP vector generally displayed levels of astrocytosis, microglial activation and storage material accumulation similar to untreated *Ppt1^-/-^* mice (Supplemental Figure 6C). Although some unexpected treatment effects of AAV9-GFP were evident in some brain regions, residual pathology was still significantly more severe in AAV9-GFP-treated *Ppt1^-/-^* mice than in WT mice (Supplemental Figure 6C).

**Figure 6.**
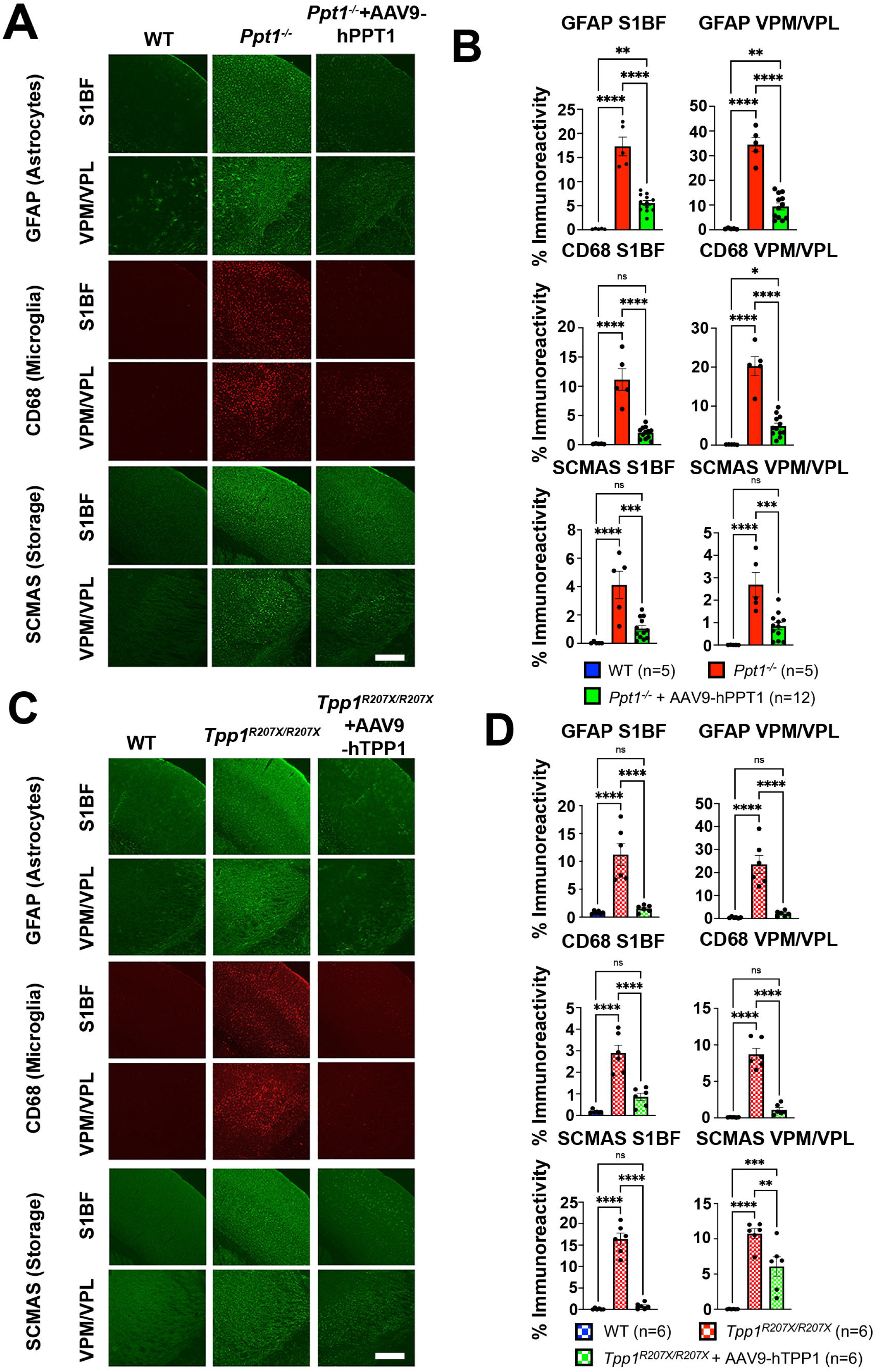
AAV9-mediated gene therapy partly attenuates neuropathological changes in the brains of *Ppt1^-/-^* and *Tpp1^R207X/R207X^* mice. **(A)** Immunostaining for subunit c of mitochondrial ATP synthase (SCMAS, green), microglial marker cluster of differentiation 68 (CD68, red), and astrocyte marker glial fibrillary acidic protein (GFAP, green) reveal the effects of neonatal AAV9-hPPT1 treatment in the brains of *Ppt1^-/-^* mice at 7 months. Scale bar 200µm **(B)** Thresholding image analysis reveals the significant treatment effects of AAV9-hPPT1 treatment upon these markers in both S1BF and VPM/VPL of *Ppt1^-/-^*mice. **(C)** Immunostaining for SCMAS (green), CD68 (red), and GFAP reveal the effects of neonatal AAV9-hTPP1- treatment upon these disease-associated neuropathological markers in the brains of *Tpp1^R207X/R207X^* mice at 3.5 months. Scale bar 200µm **(D)** Thresholding image analysis of immunoreactivity for these markers reveals the significant treatment effects of AAV9-hTPP1- treatment in both S1BF and VPM/VPL of *Tpp1^R207X/R207X^* mice. One-way ANOVA with a post-hoc Bonferroni correction (B, D), ** p≤0.01, *** p≤0.001, **** p≤0.0001.

## Discussion

Debilitating bowel dysfunction markedly impairs the quality-of-life for children with CNS neurodegenerative disorders and for their families^7, 8, 52, 53^. Understanding the basis for bowel symptoms in these disorders and developing treatments to prevent or reverse these symptoms would be a major advance, complementing new CNS-targeted treatments. Here we show for the first time that loss of lysosomal enzymes PPT1 or TPP1 causes striking and progressive damage to the myenteric plexus in CLN1 and CLN2 mice. AAV-mediated gene therapy can prevent both functional and pathological phenotypes of ENS disease. To the best of our knowledge, this is the first successful treatment of any form of enteric nervous system disease via viral-mediated gene therapy. This work may therefore have implications for the treatment of other ENS disorders or CNS neurodegenerative diseases.

Before the genetic causes of NCLs were determined, storage material in enteric neurons was used to diagnose these disorders^54^, was described in human autopsy studies^22, 23, 55–57^, and reported in animal models of CLN8 disease^58, 59^ and MPS IIIB^60^. The first decription of the LSD Niemann-Pick disease^61^ also briefly describes storage in enteric neurons, which was later confirmed in Npc mice^62^. These Npc mice did not have enteric neuron loss and only a moderate bowel transit defect^62^. Other than these few studies, ENS pathology and bowel dysfunction in lysosomal diseases appears almost entirely unexplored.

Our data suggest that the ENS forms normally in CLN1 and CLN2 disease mice. Enteric neuron loss and functional defects occur after weaning, in parallel with progressive CNS dysfunction and neurodegeneration^27, 28, 30, 41^. Interestingly, myenteric neuron loss was not uniform. Instead small regions had extensive neuron loss and other nearby areas had intact-appearing myenteric plexus, with areas in between displaying morphologically abnormal enteric neurons. These observations suggest the effects of disease within the ENS may spread from cell to cell with damaged cells changing the local environment to affect nearby cells. Consistent with this hypothesis, bowel macrophages, which depend on enteric neuron-derived colony stimulating factors 1 for survival^63^, appeared fewer in number, a phenotype plausibly linked to the loss of enteric neurons. Enteric glia also appeared affected by both PPT1- and TPP1-deficiency, with patches of reduced GFAP immunoreactivity in areas where enteric neuron loss was most pronounced, with relatively little change in the number of S100B+ enteric glia in most bowel regions. GFAP expression appeared down-regulated by enteric glia in areas where neuron loss occurs, but this marker is retained in areas where fewer neurons are lost. This is in contrast to the CNS of these mice where upregulation of both astrocyte and microglial markers occurs in areas where neuron loss is most pronounced^28, 30, 41–43^. Such differences suggest that cellular mechanisms underlying degeneration within the bowel and brain may differ.

Our data provide the first demonstration that CLN1 and CLN2 disease cause bowel-intrinsic damage to enteric neurons and glia, which may underlie the debilitating gastrointestinal symptoms experienced by children with these disorders. In support of this hypothesis, bowel contractility in *Ppt1^-/-^* mice is abnormal even when the bowel is separated from the CNS in an oxygenated organ bath. Our anatomic analyses focused on myenteric plexus neurons that coordinate muscle contraction and relaxation needed for normal bowel motility^9–13^. The observed myenteric plexus neuropathology and defects in bowel motility in oxygenated organ baths suggest that ENS neurodegeneration is a major contributor to bowel dysfunction (constipation and/or diarrhea, reflux, feeding intolerance, distension) which is common in children with NCLs. Nonetheless, CNS and peripheral nervous system pathology outside the bowel are likely to contribute to bowel dysfunction *in vivo* since sympathetic, parasympathetic, and dorsal root ganglion neurons impact bowel motility^9, 10, 64^.

Our observation that CLN1 and CLN2 disease damage the ENS has important implications for treating these disorders. Specifically, the progressive ENS damage suggests that even if CNS disease were completely prevented by gene therapy^27, 30^ or enzyme replacement therapy (ERT)^31^, ENS dysfunction would eventually cause death, unless also treated. This is a major paradigm shift since therapeutic strategies to treat NCLs predominantly target CNS disease. While these CNS-directed experimental therapies have shown promise preclinically^27, 30, 31^, and an approved CNS ERT therapy exists for CLN2 disease^65^, none are curative and none have been shown to impact the bowel.

Because lysosomal enzymes are secreted and can be taken up to ‘cross-correct’ deficient cells^49^, fewer cells need transducing for effective treatment in CLN1 or CLN2 disease than in disorders caused by deficiencies in transmembrane, mitochondrial, or cytoplasmic proteins. Remarkably, we saw pronounced treatment effects following transduction of only a small proportion of enteric neurons. More complete correction might be achieved by newer capsid variants with higher transduction efficency^66, 67^, including liver-detargeting capsids that avoid serious potential risks like hepatocarcinoma associated with intravenous AAV delivery^68^.

Because the blood brain barrier has not fully formed in neonatal mice^50, 51^, intravenous gene therapy also treated the CNS of our mice. However, IV-delivered gene therapy was less effective than our direct AAV treatment of the CNS^27, 30^, suggesting that combined intravenous and CNS-targeted delivery will be needed to improve therapeutic outcomes including survival in CLN1 and CLN2 mice.

As the first demonstration that gene therapy can successfully treat an enteric nervous system disorder in mice, our findings have direct translational consequences for the treatment of other human disorders in which ENS integrity is compromised. For ENS treatment in NCLs to become a reality, methods to effectively target the human ENS with gene therapy will need to be developed, along with detailed natural history information regarding gastrointestinal manifestations of disease. Nevertheless, our findings raise the possibility that gene therapy could treat previously underappreciated consequences of ENS disease in childhood neurodegenerative disorders.

## Abbreviations

AAV9: Adeno-associated virus 9
AFSM: Autofluorescent storage material
ANOVA: Analysis of variants
CLN1 disease: Neuronal ceroid lipofuscinosis 1
CLN2 disease: Neuronal ceroid lipofuscinosis 2
CLN8 disease: Neuronal ceroid lipofuscinosis 8
CNS: Central nervous system
DAPI: 4′,6-diamidino-2-phenylindole
ENS: Enteric nervous system
FITC: Fluorescein isothioncyanate
GFAP: Glial fibrillary acidic protein
GI: Gastrointestinal
IAUCC: Institutional Animal Care and Use Committee
LFC: Low frequency contraction
NCL: Neuronal ceroid lipofuscinoses
Npc: Niemann-Pick Type C
PPT1: Palmitoyl protein thioesterase-1
SCMAS: Subunit c of mitochondrial ATP synthase
TBS: Tris buffered saline
TBST: Tris buffered saline with Triton X-100
TPP1: Tripeptidyl peptidase-1
TTX: Tetrodotoxin
vg: Viral genomes

## Acknowledgments

We thank Dr. Jill M. Weimer and Dr. David A. Pearce (Sanford Research) for supplying the *Cln2^R207X^* mice that were used in this study. We also thank the Washington University Center for Cellular Imaging (WUCCI) supported by Washington University School of Medicine, the Children’s Discovery Institute of Washington University and St. Louis Children’s Hospital, and the Foundation for Barnes-Jewish Hospital in St. Louis, for providing a Zeiss Axio Scan Z1 Fluorescence Slide Scanner. We also acknowledge Drs. Patricia Dickson, Phillip Tarr, Ewa Ziółkowska and Alison Barnwell for constructive comments on the manuscript.

## Study Approval

All animal procedures were performed in accordance with National Institutes of Health (NIH) guidelines under protocols 2018-0215 and 21-0292 approved by the Institutional Animal Care and Use Committee (IACUC) at Washington University School of Medicine in St. Louis, MO; and The Children’s Hospital of Philadelphia IACUC protocol IAC 22-001041.

---

**Supplemental Table 1:** List of antibodies used

| **Antibody** | **Concentration** | **Catalog number** | **Source** |
| --- | --- | --- | --- |
| ANNA-1 (HuC/D) | 1:10000 (bowel) | N/A | Kind gift from Dr. Vanda Lenon, Mayo Clinic; RRID: AB_2314657 |
| Rabbit anti-GFAP | 1:10000 (bowel)<br>1:1000 (brain) | Z0334 | Agilent<br>RRID: AB_10013382 |
| Rabbit anti-beta III tubulin (Tuj1) | 1:5000 (bowel) | ab18207 | Abcam<br>RRID: AB_444319 |
| Rabbit anti-Iba1 | 1:2000 (bowel) | 019-19741 | FUJIFILM Wako<br>RRID: AB_839504 |
| Rat anti-CD68 | 1:400 (brain) | MCA1957 | Biorad<br>RRID: AB_322219 |
| Rabbit anti-S100 beta | 1:300 (bowel) | ab52642 | Abcam<br>RRID: AB_882426 |
| Rabbit anti-SCMAS | 1:400 (brain) | ab181243 | Abcam<br>RRID: AB_2935765 |
| AlexaFluor goat anti-human 546 | 1:400 (bowel & brain) | A-21089 | ThermoFisher Scientific (Invitrogen)<br>RRID: AB_2535745 |
| AlexaFluor goat anti-rat 546 | 1:400 (brain) | A-11081 | ThermoFisher Scientific (Invitrogen)<br>RRID: AB_2534125 |
| AlexaFluor goat anti-rabbit 488 | 1:400 (bowel & brain) | A-11008 | ThermoFisher Scientific (Invitrogen)<br>RRID: AB_143165 |

**Supplemental Figure 1: Bowel contractility kymographs in *Ppt1^-/-^* mice in the presence and absence of tetrodotoxin**. Bowel contractility kymographs were recorded in small intestine isolated from endstage *Ppt1^-/-^* mice and wild type controls at 7 months. There were markedly fewer and disorganised low frequency contractions (LFCs) (white arrowheads on kymographs) in *Ppt1^-/-^* mice (quantification in Figure 1F). These LFCs were abolished by tetrodotoxin administration, suggesting they are mediated by enteric nervous system activity.

**Supplemental Figure 2: Lack of enteric pathology in NCL mouse models at 1 month of age**. (A) Merged photomicrographs showing immunostaining for HuC/D (neurons, red) and the glial marker S100B (green) in the bowel of wild type (WT), *Ppt1^-/-^* and *Tpp1^R207X/R207X^* mice at 1 month of age reveal no loss of either neurons or glia at this age. Scale bar 200µm. (B) Counts of the density of HuC/D positive neurons and S100B positive enteric glia reveal no significant difference in any bowel regions of 1 month old *Ppt1^-/-^* mice, *Tpp1^R207X/R207X^* mice or age matched wildtype (WT controls).

**Supplemental Figure 3**: **Evidence for enteric nervous system pathology in *Ppt1^-/-^* mice.** Photomicrographs showing immunostaining for HuC/D (neurons, red) and and either the neurofilament marker Tuj1 (green), the glial marker glial fibrillary acidic protein (GFAP, green), the glial marker S100B (S100B, green), or the macrophage marker Iba1 (green) in the duodenum, ileum and colon of *Ppt1^-/-^* mice and age-matched wildtype (WT) controls at 7 months of age. (Merged channel images of these figures are included in Figure 2D). Scale bar 200µm.

**Supplemental Figure 4**: **Evidence for enteric nervous system pathology in *Tpp1^R207X/R207X^*mice.** Photomicrographs showing immunostaining for HuC/D (neurons, red) and either the neurofilament marker Tuj1 (green), the glial marker glial fibrillary acidic protein (GFAP, green), the glial marker S100B (S100B, green), or the macrophage marker Iba1 (green) in the duodenum, ileum and colon of *Tpp1^R207X/R207X^* mice and age-matched wildtype (WT) controls at 3.5 months. (Merged channel images of these figures are included in Figure 3C). Scale bar 200µm.

**Supplemental Figure 5: Increased soma size of Iba1 positive macrophages in *Ppt1^-/-^* and *Tpp1^R207X/R207X^* mice. (A)** Histograms reveal the significantly increased cell soma size of Iba1 positive macrophages in bowel of disease end stage *Ppt1^-/-^* mice (7 months) and *Tpp1^R207X/R207X^*mice (3.5 months), compared to age-matched wild type (WT) controls. **(B)** Cell size distribution graphs reveal the shift to larger macrophage soma size in all bowel regions of *Ppt1^-/-^* mice (7 months) and *Tpp1^R207X/R207X^* mice (3.5 months), compared to age-match WT controls. Two tailed *t* test, **** p≤0.0001.

**Supplemental Figure 6: Effects of gene therapy with AAV9-GFP. (A)** Immunostained bowel wholemount preparations reveal the transduction of HuC/D positive (red) myenteric neurons by AAV9-GFP (green, resulting in yellow labelling of transduced neurons in this merged image) in AAV9-GFP-treated *Ppt1^-/-^* mice, but not in age-matched untreated *Ppt1^-/-^* or wild type (WT) mice. Scale bar 100µm. **(B)** Counts of the number of HuC/D positive neurons reveal the lack of therapeutic effect in any bowel region of AAV9-GFP-treated *Ppt1^-/-^*mice, compared to the significant treatment effect of AAV9-hPPT1-treated *Ppt1^-/-^*mice (see Figure 4). **(C)** Effects of intravenous neonatal administration of AAV9-GFP in the brains of *Ppt1^-/-^* mice. Compared to untreated *Ppt1^-/-^*mice, AAV9-GFP-treated *Ppt1^-/-^* mice show no obvious treatment effects upon astrocytosis (GFAP, green), microglial activation (CD68, red) or storage material accumulation (SCMAS, green), as is evident in photomicrographs and in thresholding image analysis. The exception is in the ventral posterior thalamic nucleus (VPM/VPL) where significant treatment effects of AAV9-GFP were evident upon astrocytosis and microglial activation, but not storage material accumulation. No treatment effects were evident in the primary somatosensory cortex (S1BF). Scale bar 200µm. One-way ANOVA with a post-hoc Bonferroni correction (B, C), * p≤0.05; ** p≤0.01, **** p≤0.0001.

