## Supplementary figures and images for "Bowel dysmotility and enteric neuron degeneration in lysosomal storage disease mice is prevented by gene therapy"

### Supplemental Figures

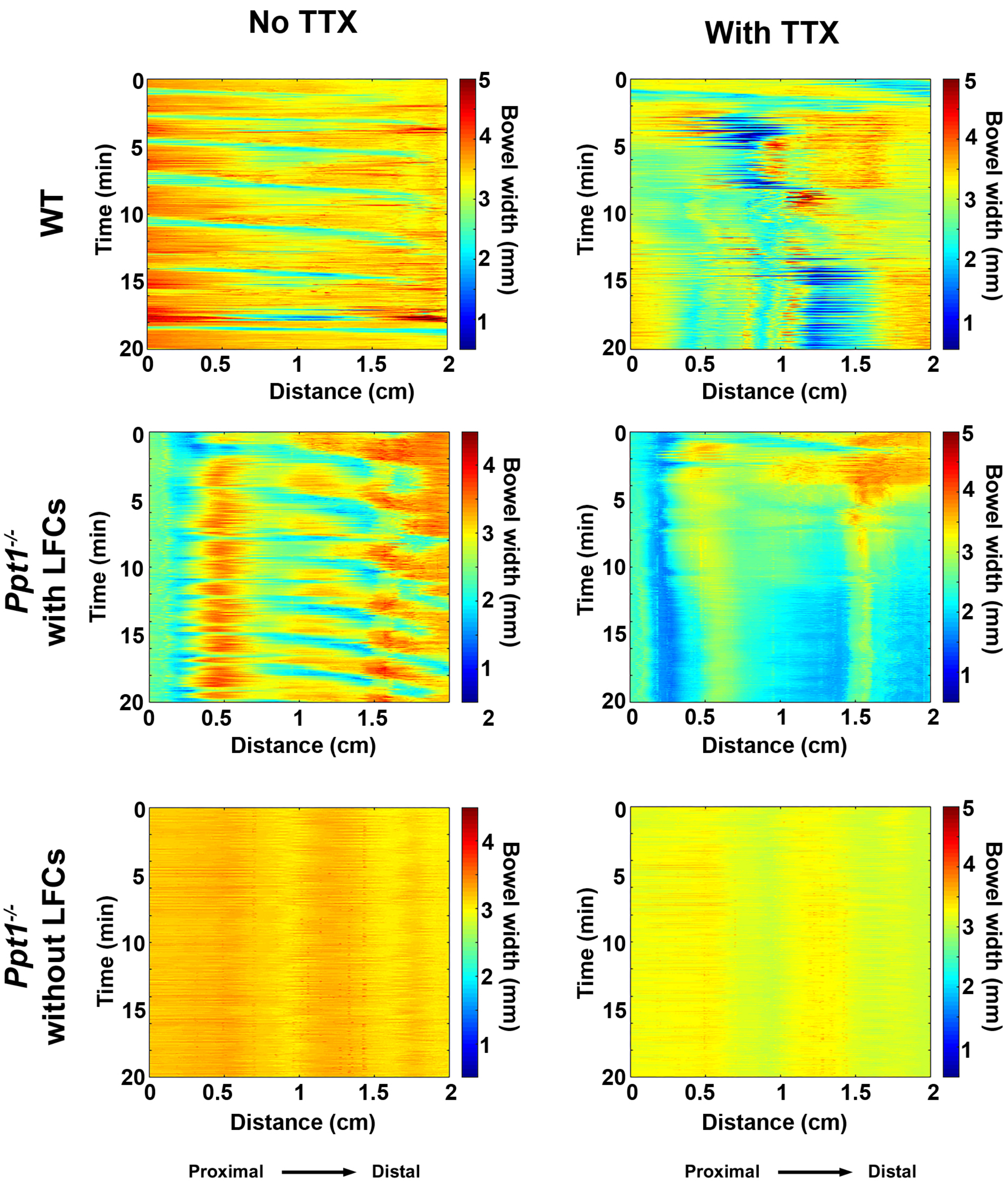

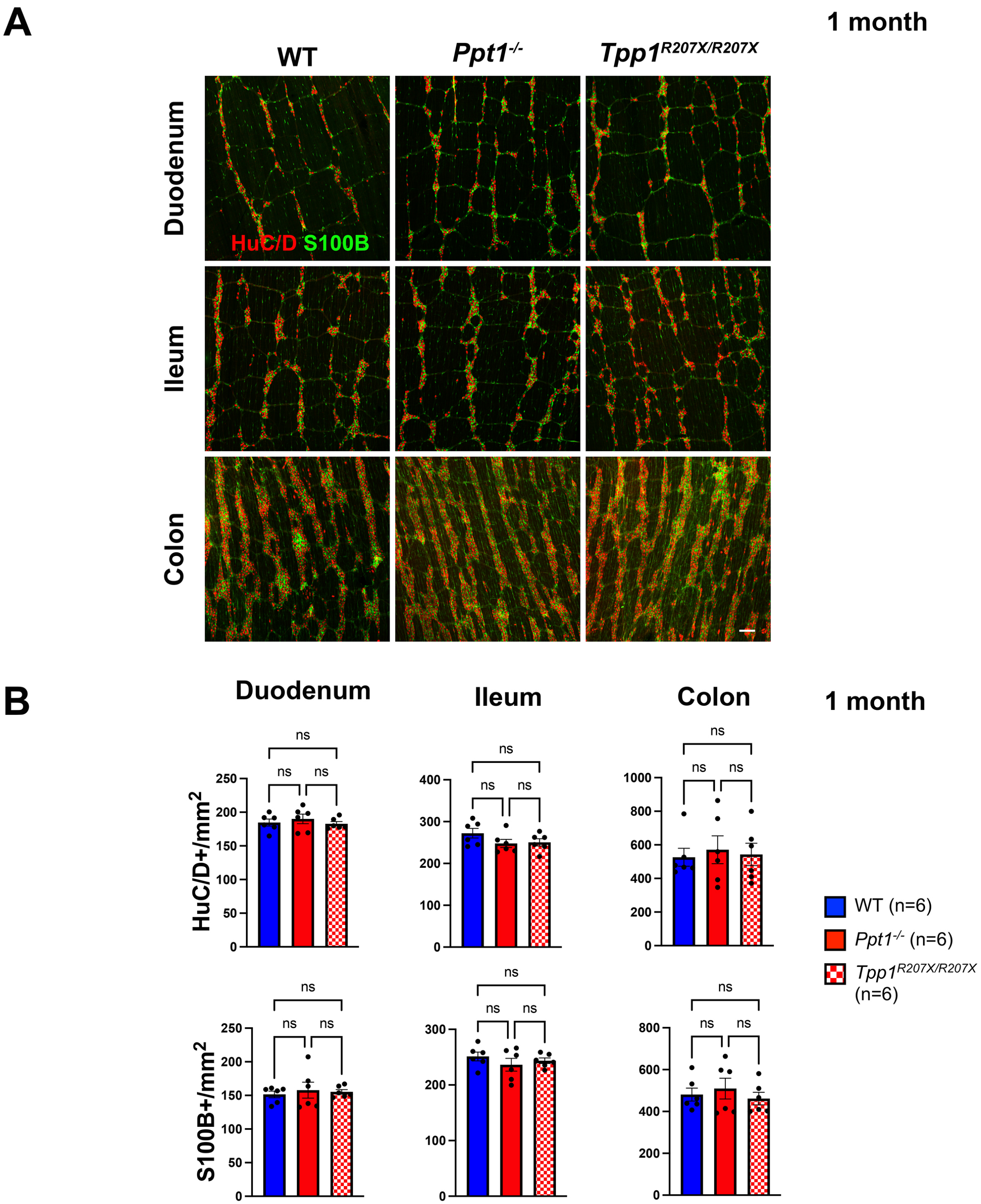

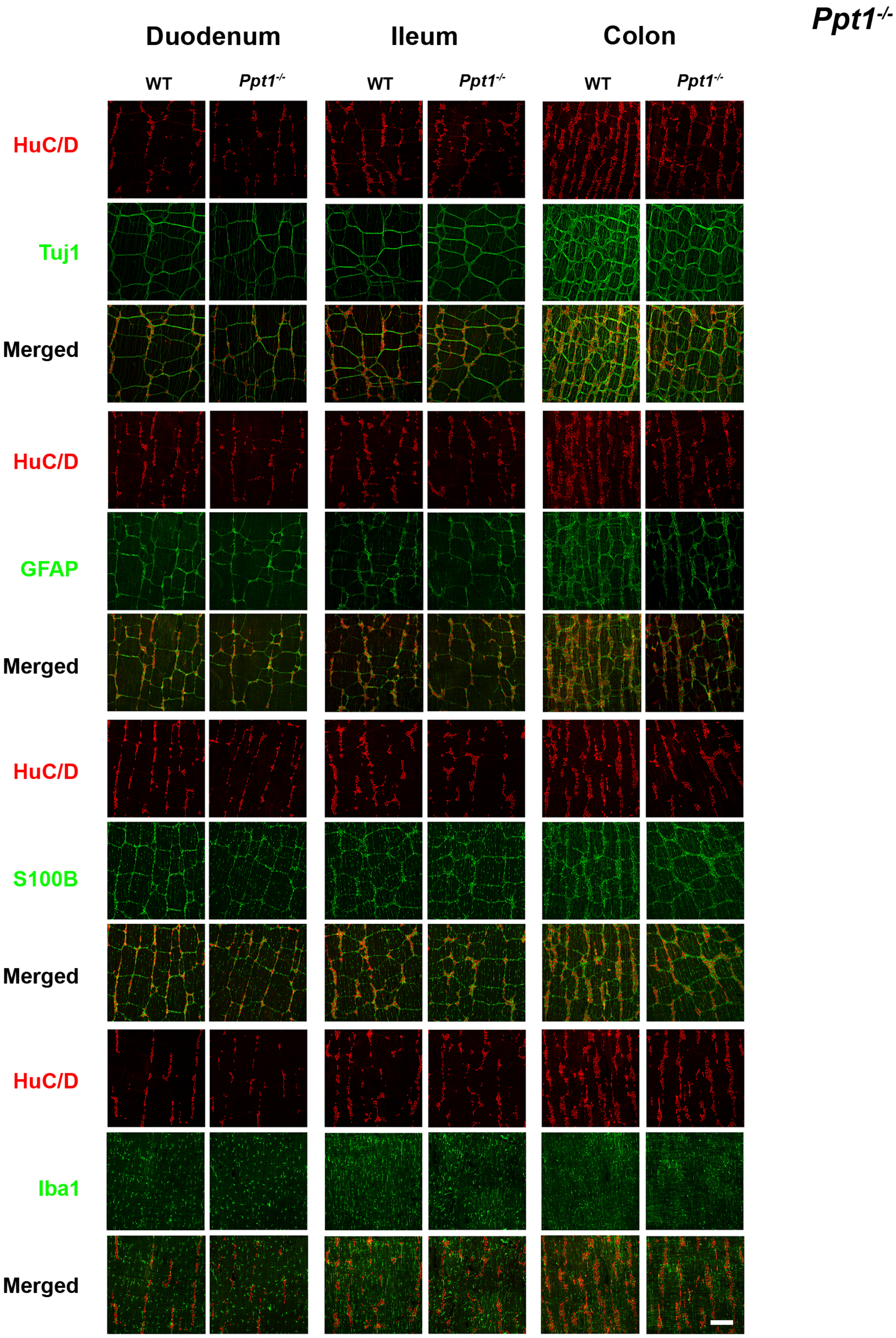

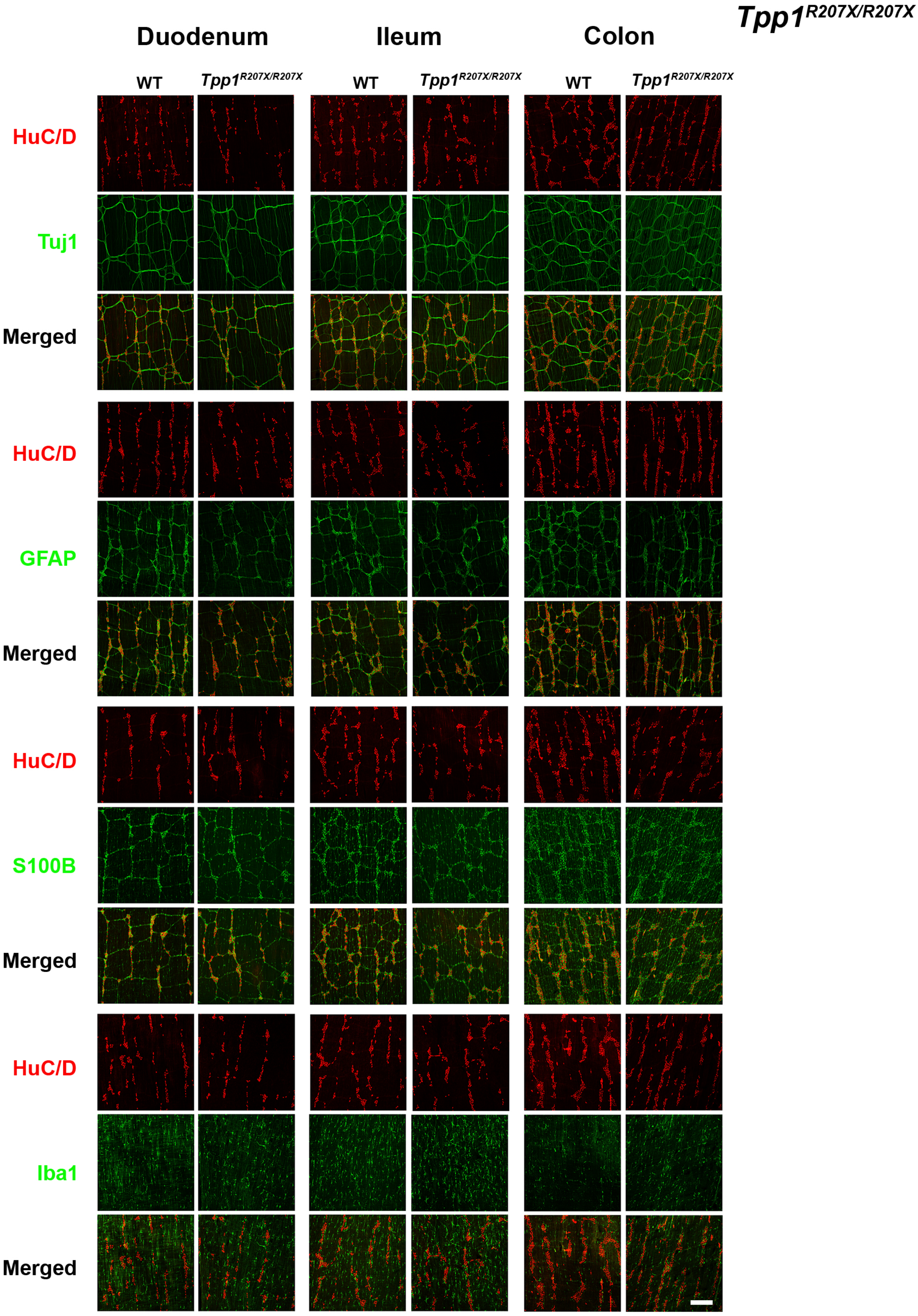

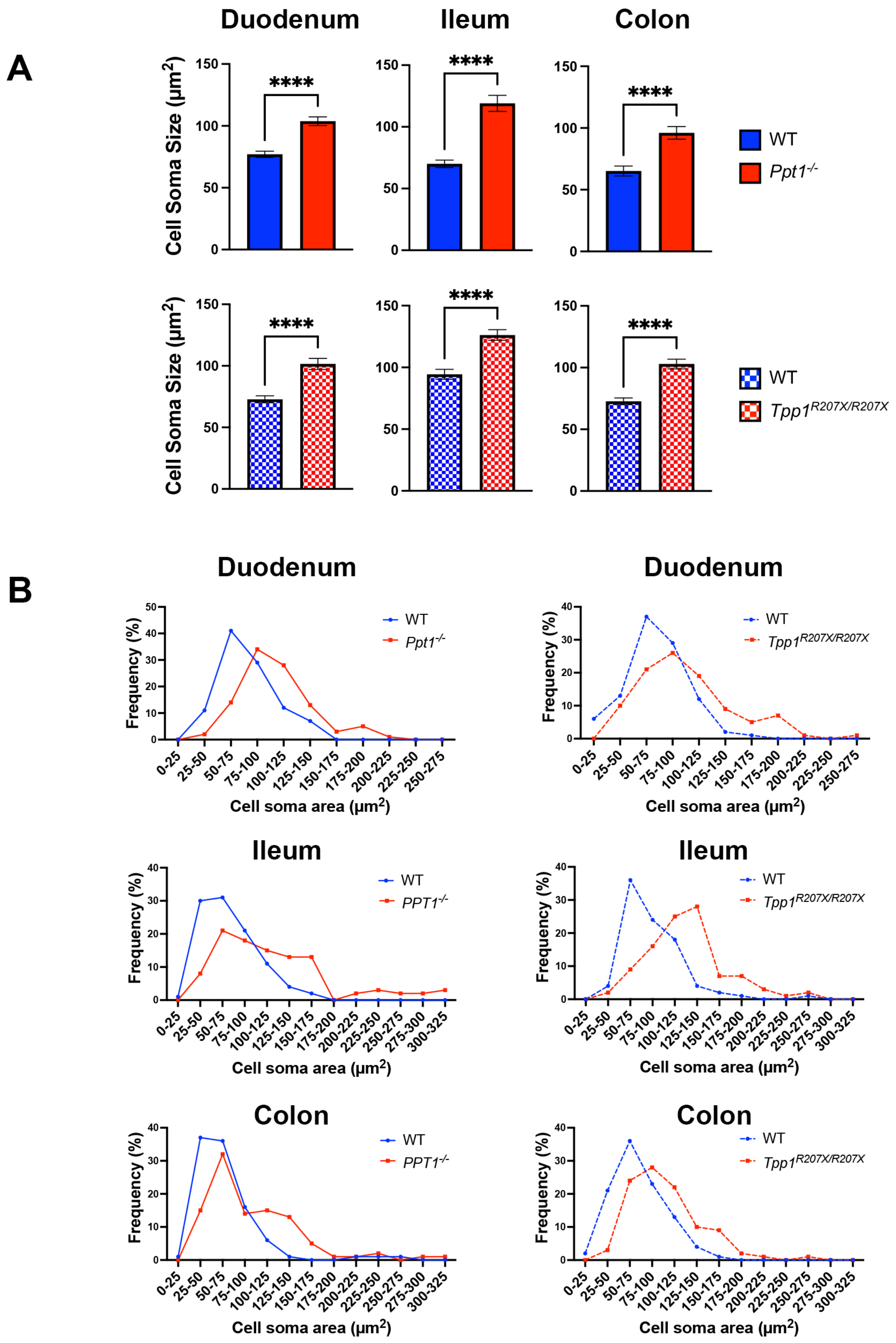

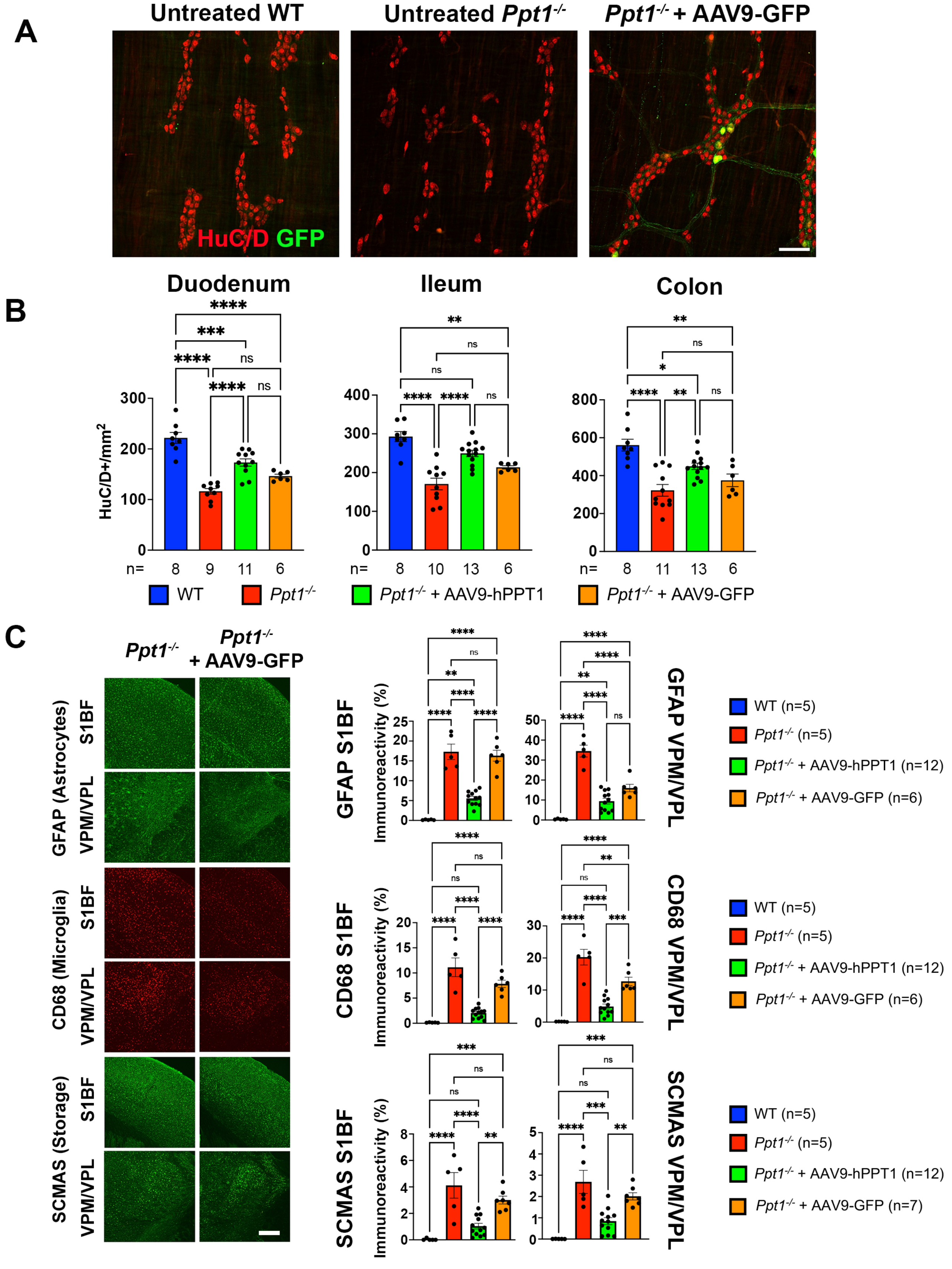
